# Acyloxyacyl Hydrolase is a Host Determinant of Gut Microbiome-Mediated Pelvic Pain

**DOI:** 10.1101/2021.01.27.428290

**Authors:** Afrida Rahman-Enyart, Wenbin Yang, Ryan E. Yaggie, Bryan White, Michael Welge, Loretta Auvil, Matthew Berry, Colleen Bushell, John M. Rosen, Charles N. Rudick, Anthony J. Schaeffer, David J. Klumpp

**Author notes:** Address all correspondence to, 16-719 Tarry Building, 303 East Chicago Avenue, Chicago, IL, P, F. These authors contributed equally (Afrida Rahman-Enyart, Wenbin Yang).

## Abstract

Dysbiosis of gut microbiota is associated with many pathologies, yet host factors modulating microbiota remain unclear. Interstitial cystitis/bladder pain syndrome (IC/BPS or “IC”) is a debilitating condition of chronic pelvic pain often with co-morbid urinary dysfunction and anxiety/depression, and recent studies find fecal dysbiosis in IC/BPS patients. We previously identified the locus encoding acyloxyacyl hydrolase, *Aoah*, as a modulator of pelvic pain severity in a murine IC/BPS model. AOAH-deficient mice spontaneously develop rodent correlates of pelvic pain, increased responses to induced pelvic pain models, voiding dysfunction, and anxious/depressive behaviors. Here, we report that AOAH-deficient mice exhibit dysbiosis of GI microbiota. AOAH-deficient mice exhibit an enlarged cecum, a phenotype long associated with germ-free rodents, and reduced trans-epithelial electrical resistance consistent with a “leaky gut” phenotype. AOAH-deficient ceca showed altered gene expression consistent with inflammation, Wnt signaling, and urologic disease. 16S rRNA sequencing of stool revealed altered microbiota in AOAH-deficient mice, and GC-MS identified altered metabolomes. Co-housing AOAH-deficient mice with wild type mice resulted in converged microbiota and altered predicted metagenomes. Co-housing also abrogated the pelvic pain phenotype of AOAH-deficient mice, which was corroborated by oral gavage of AOAH-deficient mice with stool slurry of wild type mice. Converged microbiota also alleviated comorbid anxiety-like behavior in AOAH-deficient mice. Oral gavage of AOAH-deficient mice with anaerobes cultured from IC/BPS stool resulted in exacerbation of pelvic allodynia. Together, these data indicate that AOAH is a host determinant of normal gut microbiota, and the dysbiosis associated with AOAH deficiency contributes to pelvic pain. These findings suggest that the gut microbiome is a potential therapeutic target for IC/BPS.

## INTRODUCTION

Interstitial cystitis/bladder pain syndrome (IC/BPS or “IC”) is a chronic condition characterized by bladder discomfort and moderate to severe pelvic pain affecting as many as eight million patients in the United States [1–3]. Patients suffering from IC/BPS commonly exhibit co-morbid conditions including voiding dysfunction of increased urinary frequency and urgency and anxiety/depression [4–6]. The etiology of IC/BPS is unknown and pathophysiologic mechanisms remain unclear, so IC/BPS is a clinical challenge with no widely effective therapy. Because of the profound impact of chronic pelvic pain, the National Institute of Diabetes and Digestive and Kidney Diseases (NIDDK) established the Multi-Disciplinary Approaches to Chronic Pelvic Pain (MAPP) Research Network in an effort to study urologic chronic pelvic pain syndromes at the clinical, epidemiologic, and basic levels. In our MAPP studies, we previously conducted a genetic screen for loci modulating pelvic pain severity and identified *Aoah*, the locus encoding acyloxyacyl hydrolase, as a modulator of pelvic pain in a murine model of IC/BPS [7].

AOAH is a host enzyme best known for detoxifying bacterial lipopolysaccharides (LPS) by selectively removing secondary acyl chains from the lipid A moiety, mitigating the host response to bacterial infection and attenuating sustained inflammation [8–12]. Previous studies in our lab demonstrated that mice deficient for AOAH spontaneously develop pelvic allodynia consistent with pelvic pain and exhibit heightened response in induced pelvic pain models [7]. AOAH-deficient mice also have voiding dysfunction [13], and display rodent correlates of anxious/depressive behaviors [14]. Expressed along the bladder-brain axis, AOAH functions as an arachidonyl acyl transferase, and AOAH-deficient mice exhibit defects in central nervous system (CNS) arachidonic acid homeostasis and increased accumulation of PGE2, contributing to pelvic pain [15]. Moreover, corticotropin-releasing factor (CRF) is the initiator of the hypothalamic-pituitary-adrenal (HPA) axis and a modulator of pain responses, and AOAH-deficient mice have increased *Crf* expression and increased serum corticosterone consistent with HPA axis dysregulation [14]. IC/BPS patients also have aberrant diurnal cortisol fluctuations indicating HPA axis dysregulation [16], another similarity with AOAH-deficient mice. Thus, AOAH-deficient mice recapitulate several key aspects of IC/BPS.

Gut dysbiosis has been increasingly recognized as a key player in visceral pain, and recent studies have indicated altered fecal microbiota in women with IC/BPS as well as in male patients suffering an analogic condition, chronic prostatitis/chronic pelvic pain syndrome [17–20]. Visceral convergence of sensory inputs that drives pelvic organ crosstalk and the gut-brain axis provide mechanisms by which gut dysbiosis may influence IC/BPS urologic and cognitive symptoms [21–25]. Given the strong parallels between phenotypes of AOAH-deficient mice and IC/BPS, we hypothesized that AOAH-deficient mice would exhibit gut dysbiosis that mediates pelvic pain and other comorbidities. We observed multiple phenotypes supporting this hypothesis including altered cecal morphology and transcriptomes, fecal and cecal dysbiosis, and a role for gut microbiota in pelvic pain behaviors.

## MATERIALS AND METHODS

### Animals

Ten to twelve-week-old male and female wild type (WT) C57BL/6 mice were purchased from The Jackson Laboratory. *Aoah*^—/—^ mice (B6.129S6-Aoahtm1Rsm/J) were obtained as a generous gift from Dr. Robert Munford of NIAID and maintained on a 12h:12h light:dark cycle as previously described [14]. All animals were maintained at the Center for Comparative Medicine under Northwestern IACUC approved protocols.

### GI characterization

Animals were euthanized and placed on a pan balance to obtain body weight. The cecum and bladder were excised and weighed. The cecum was then dissected, and cecal stool was removed and weighed.

### Transepithelial electrical resistance

Cecum and bladder permeability were measured as transepithelial electrical resistance (TEER) as previously described for bladders in Aguiniga et al., 2020 [13]. Excised organs were bisected and immediately placed in Ringer’s solution at room temperature. After rinsing, ceca and bladders were mounted onto cassettes with apertures of 0.126cm^2^ and 0.04cm^2^ respectively. Cassettes were inserted between two-halves of an Ussing chamber (Physiologic Instruments EM-CSYS-2). KCl saturated salt-bridge electrodes were inserted. The chambers were filled with Ringer’s solution and bubbled with carbogen (95% O_2_/5% CO_2_) and maintained at 37 °C until preliminary resistance became stabilized, typically at least 1 hr. Voltage was clamped and current passed every 3 s at 30 s intervals using a VCC MC2 multichannel voltage-current clamp amplifier controlled with Acquire and Analyze software v2.3.4 (Physiologic Instruments).

### Hematoxylin and eosin staining

All hematoxylin and eosin (H&E) staining in wild type and AOAH-deficient ceca was performed by Northwestern University Mouse Histology & Phenotyping Laboratory (MHPL).

### Cecum staining

Animals were euthanized and ceca were dissected. The cecum was flushed first with PBS and then 10% Neutral Buffered Formalin (NBF). Tissue was Swiss-rolled and placed in 15 ml of 10% NBF prior to embedding in paraffin. Tissue was then sectioned at 4 μm. Tissue was de-paraffinized with a series of washes as follows: 2 × 3 min with 100% xylene, 3 × 3 min with 100% ethanol, 1 × 3 min with 94% ethanol, 1 × 3 min with 70% ethanol, 1 × 3 min with 50% ethanol, and 1 × 3 min with H_2_O. Antigen retrieval was performed in a pressure cooker using retrieval solution (DAKO, Santa Clara, CA) at 100 °C for 10 min, 90 °C for 10 min, and then washed in PBS for 3 min. Tissue was blocked in FC Receptor Block (Innovex Biosciences, Richmond, CA) for 10 min and Background Buster (Innovex) for 30 min. Tissue was incubated with AOAH primary antibody (Abcam, Cambridge, United Kingdom) at 1:100 in Antibody Dilution Buffer (DAKO) overnight at 4 °C. Tissue was then washed in 1X PBS 3 × 5 min followed by incubation with Alexa Fluor 488 secondary antibody (Invitrogen, Carlsbad, CA) diluted to 1:500 in Antibody Dilution Buffer (DAKO) for 2 hrs at room temperature. Secondary was washed in 1X PBS 3 × 5 min and nuclei were stained using DAPI followed by imaging.

### Gut microbiome analyses

Fecal and cecal gut microbiota were analyzed as previously reported in Braundmeier-Fleming et al., 2016 [19]. Briefly, DNA was extracted from fecal pellets or cecal stool by QIAamp DNA Stool Mini Kit (QIAGEN, Hilden, Germany) by homogenizing with a Mini-Beadbeater and 0.1 mm zirconia/silica beads (Biospec, Bartlesville, OK) in 1.4 mL ASL buffer (QIAGEN). Phylotype profiles of microbiota were generated by deep amplicon sequencing of the 16S rRNA V3–V5 hypervariable region. Barcoding samples followed by MiSeq tag sequencing yielded approximately 50,000 reads/sample, detecting dominant and poorly represented taxa. The V3–V5 hypervariable region was selectively amplified through 30 cycles with primers 357F (CCTACGGGAGGCAGCAG) and 926R (CCGTCAATTCMTTTRAGT). Amplicon pools were quantified on a Qubit fluorimeter, and fragment sizes were determined by Agilent bioanalyzer High Sensitivity DNA LabChip (Agilent Technologies, Wilmington, DE) followed by dilution to 10 nM. To accurately calculate matrix, phasing, and prephasing, amplicons were spiked with a PhiX control library to 20%. Mixtures were sequenced on an Illumina MiSeq V2 (250nt from each end). To define OTU abundance profiles and phylogenic relationships, sequence reads were binned at 97% identity using QIIME and Galaxy.

### Predictive functional analysis

Phylogenetic Investigation of Communities by Reconstruction of Unobserved States (PICRUSt) was implemented using 16S rRNA sequence data from fecal samples to predict function as previously described [19, 26].

### Fecal metabolomics

Analyses of fecal and cecal gut metabolomes were analyzed as previously reported in Braundmeier-Fleming et al., 2016 [19]. Briefly, duplicate samples were dried, and one was derivatized [27] with minor modifications: 90 minutes at 500 °C with 80 μl of methoxyamine hydrochloride in pyridine (20 mg/ml) followed by incubating 60 min at 500 °C with 80 μl MSTFA. A 5 μl aliquot of a C31 fatty acid internal standard was added to each; in derivatized samples this occurred prior to trimethylsilylation. 1 ml samples were injected with a split ratio of 7:1 into an Agilent GC-MS system (Agilent Inc, Palo Alto, CA) consisting of an 7890A gas chromatograph, a 5975C mass-selective detector, and a 7683B autosampler. GC was performed on 60 m HP-5MS columns (0.25 mm inner diameter, 0.25 mm film thickness; Agilent Inc) at 2500 °C injection and interface temperatures and 2300 °C ion source and helium carrier flow rate of 1.5 ml min^−1^. A 5-min isothermal heating program at 700 °C was followed by a 50 °C min^−1^increase to 3100 °C and finally 20 min at 3100 °C. The mass spectrometer was operated in positive electron impact mode at 69.9 eV ionization energy in a scan range of m/z 30–800. All resulting chromatogram were compared with mass spectrum libraries NIST08 (NIST, Gaithersburg, MD), WILEY08 (Palisade Corporation, Ithaca, NY), and a University of Illinois Metabolomics Center custom library. All data were normalized to internal standards to facilitate direct comparisons, and chromatograms and mass spectra were evaluated using MSD ChemStation (Agilent) and AMDIS (NIST), where retention time and mass spectra were implemented within the AMDIS method formats.

### Cecum RNA preparation and microarray

The cecum was dissected from mice immediately following euthanasia by cervical dislocation and stored at −80°C. Cecal epithelium was homogenized in ice-cold TRIzol with a homogenizer, and total RNA was purified according to manufacturer’s instructions (Invitrogen, Carlsbad, CA). Gene expression was quantified using Affymetrix Mouse Genome 430 2.0 array that contain 45,101 probes and measure the expression level of 20,022 unique NCBI Entrez-identified genes. The data sets were preprocessed with normalization, variance stabilization, and log_2_ transformation. Student’s *t*-tests were used to identify genes significantly differentially expressed (*P* < 0.05 and 2-fold) between WT and AOAH-deficient mice. Average difference values were normalized to median over the arrays. The data were filtered so that only those genes that were adequately measured on 75% of the arrays were included. A class comparison protocol was used to identify genes whose degree of expression differed significantly by twofold or more among the three groups. To visualize whole genome expression level by function and pathway, the microarray data were analyzed with MetaCore Analysis (IPA; Ingenuity Systems, Redwood City, CA). IPA analysis identified canonical pathways differentially expressed (*P* < 0.05) between WT and AOAH-deficient mice.

### Co-housing and fecal microbiota transfer (FMT) experiments

For co-housing experiments, two WT and two AOAH-deficient mice were housed in the same cage (4 mice/cage) for at least 1 week prior to experimentation. For FMT experiments, mice from both groups received oral FMT from either WT or AOAH-deficient mice. Fecal suspensions were prepared from three FMT donor mice by suspending one fecal pellet in 1 mL sterile phosphate buffered saline using a syringe and 18-gauge needle (Becton Dickinson, Vernon Hills, IL) until the entire volume could pass through the needle. We allowed particulates to settle for 30 s before drawing supernatant into a syringe attached to a 22-gauge gavage needle (Fine Science Tools, Foster City, CA). We pooled together 500 μL of the supernatants from each of the three 1 mL suspensions and deposited 200 μL of this pooled supernatant into the stomach of each recipient mouse by gavage. Each FMT recipient was gavaged every other day for 7 days. We cleaned gavage needles with 70% ethanol between mice, using different gavage needles for each treatment group.

### FMT of human samples

Human stool samples from healthy patients and patients with IC/BPS were placed in pre-reduced PBS with 0.1% cysteine (PBSc). Diluted (10^−4^) samples were plated on 10cm plates and grown under anaerobic conditions at 37 °C. Once plates were dense they were scraped into PBSc. 100ul of sample was gavaged into male mice 3 × 3 days. After 7 days mice were tested for tactile allodynia to assess pelvic pain.

### Pelvic allodynia

Mice were tested for pelvic allodynia using von Frey filament stimulation to the pelvic region, as previously described in Rudick et al., 2012 [28]. Briefly, prior to co-housing, baseline pelvic allodynia was measured for WT and AOAH-deficient mice. Mice were co-housed for 28 d followed by testing. After co-housing, mice were placed in a test chamber and allowed to acclimate for 5 min. Starting with the lowest force filament, five von Frey filaments were applied 10 times to the pelvic region. A response was considered to be painful if the animal jumped, lifted and shook the hind paws, or excessively licked the pelvic region.

### Urinary bladder distention evoked visceromotor response (VMR)

Mice were anesthetized with 2% isoflurane. Silver wire electrodes were placed on the superior oblique abdominal muscle and subcutaneously across the abdominal wall (as a ground) to allow differential amplification of the abdominal VMR signals. A lubricated angiocatheter was inserted into the bladder via the urethra for bladder distention. After completion of the surgical preparation, isoflurane anesthesia was reduced to ∼0.875% until a flexion reflex response was present (evoked by pinching the paw) but spontaneous escape behavior and righting reflex were absent. The animals were not restrained in any fashion. Body temperature was maintained using an overhead radiant light and monitored throughout the experiment. The conditions were optimized to establish a stable depth of anesthesia and consistent baseline VMR activity. Phasic bladder distention with PBS was then used to evoke bladder nociception. The PBS pressure was controlled by an automated distention control device custom made in the Washington University School of Medicine Electronic Shop. The distention stimulus applied 20–80 mmHg pressure for 20 seconds every 2 min. The VMR signal was relayed in real-time using a Grass CP511 preamplifier to a PC via a WinDaq DI-720 module. Data were exported to Igor Pro 6.05 software (Wavemetrics, Portland, OR). Using a custom script, VMR signals were subtracted from the baseline, rectified and integrated over 20 s to quantify the area under the curve and presented in arbitrary units. The investigator who quantified the VMR was blinded to treatment.

### Defensive burying

In order to measure anxiety-like behavior, AOAH-deficient and WT mice were singly placed in a novel holding cage with 7 cm of fresh bedding for 15 min. Anxiety-like behavior was recorded as latency of first (measured in seconds) and number of burying behaviors. Burying behavior was defined as the mouse aggressively sifting through bedding with both front paws and with their head facing down.

### Statistical analysis

Results were analyzed by Student’s t-test or one-way/two-way analysis of variance (ANOVA) followed by Tukey’s Multiple comparisons test with the use of Prism software, version 6 (GraphPad, Inc). Differences were considered statistically significant at P<0.05.

## RESULTS

### AOAH-deficient mice have enlarged ceca and increased gut permeability

Dissection of AOAH-deficient mice suggested a GI phenotype relative to wild type mice where AOAH-deficient mice ceca appeared larger and distended in both females (Fig. 1A) and males (Fig. S1A). This difference in appearance was at least partially borne out in mass because the cecum and cecal stool as a function of body mass were significantly heavier in male AOAH-deficient mice compared to wild type mice (Fig. S1B). In contrast, despite the enlarged morphology of AOAH-deficient ceca, neither cecal tissue nor cecal stool were significantly more massive in AOAH-deficient females relative to wild type mice (Fig. 1B). We further characterized the cecum phenotype by assessing cecum barrier function in female and male wild type and AOAH-deficient mice by quantifying cecum transepithelial electrical resistance (TEER) ex vivo via an Üssing chamber (Fig. 1C and Fig. S1C respectively). Cecum TEER was significantly lower in AOAH-deficient mice compared to wild type in both females (33.58 ± 4.77 Ω•cm^2^ for wild type and 19.94 ± 6.66 Ω•cm^2^ for *Aoah*^*—/—*^; Fig. 1C) and males (27.66 ± 6.63 Ω•cm^2^ for wild type and 17.11 ± 6.11 Ω•cm^2^ for *Aoah*^*—/—*^; Fig. S1C), indicating that AOAH-deficient mice have increased cecum permeability or “leakiness” in both sexes. Since IC/BPS more commonly afflicts women, we focused further characterization on female mice.

**Figure 1.**
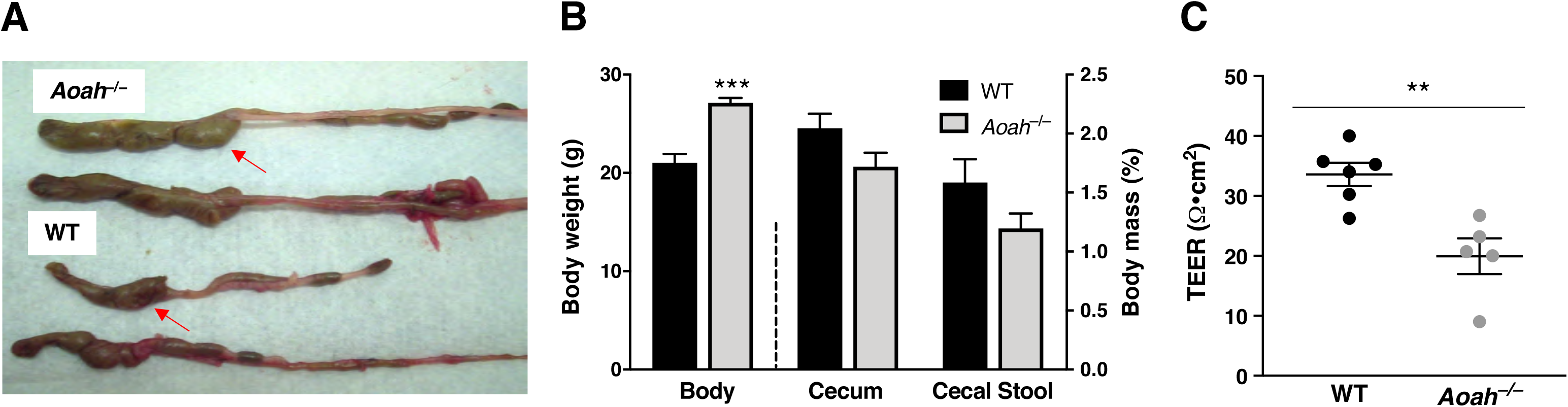
Female acyloxyacyl hydrolase-deficent (*Aoah*^—/—^) mice exhibit enlarged ceca and compromised gut permeability. **A**. Comparison of representative cecum sizes (arrows) of female wild-type (WT) C57BL/6 (bottom) and *Aoah*^—/—^ (top) mice. **B**. Female *Aoah*^—/—^ mice showed increased body weight versus WT mice (WT mice: n=4 and *Aoah*^—/—^ mice n=7, P<0.0001, Student’s t test, two tailed). No differences were observed in the mass of the cecum and cecal stool (WT mice: n=4 and *Aoah*^—/—^ mice n=6, P > 0.05, Student’s t test, two tailed). Bars represented as average ± SEM **C**. Ceca of female *Aoah*^—/—^ mice showed a decrease in transepithelial electrical resistance (TEER) compared to WT (WT mice: n=6 and *Aoah*^—/—^ mice n=5, P<0.01, Student’s t test, two tailed). Data represented as individual values and average (represented by horizontal line) ± SEM.

### AOAH-deficient mice have altered gut signaling pathways

The compromised cecal TEER of AOAH-deficient mice suggest functionally altered epithelium, yet histologic examination uncovered no obvious differences in cecal epithelia of wild type and AOAH-deficient mice (Fig. 2A, top row). Immunostaining of AOAH protein in wild type mouse cecum revealed sparse expression of AOAH in the epithelium, which was mainly absent in AOAH-deficient cecum (Fig. 2A, bottom row). Transcriptome profiling by microarray analyses revealed significant differences in gene expression consistent with altered signaling pathways and disease risk in the AOAH-deficient ceca (Fig. 2B, C, and S2 respectively). Examination of genes increased/decreased at least 2-fold indicated altered mRNA abundance in AOAH-deficient mice of genes associated with immune response/inflammation (53%), epithelia (13%), bacteria (7%), lipids (6%), neurons (5%), cell survival (5%), protein transport (4%), thioredoxin (2%), P450 (2%), and erythrocytes (2%) (Fig. 2B). Consistent with this, pathway analyses also revealed increased immune response signaling in AOAH-deficient mice compared to wild type (Fig. 2C). Together, these findings suggest that an inflammatory millieu and alterations in mRNA associated with epithelia contribute to increased cecal epithelial permeability underlying a leaky gut phenotype of AOAH-deficient mice.

**Figure 2.**
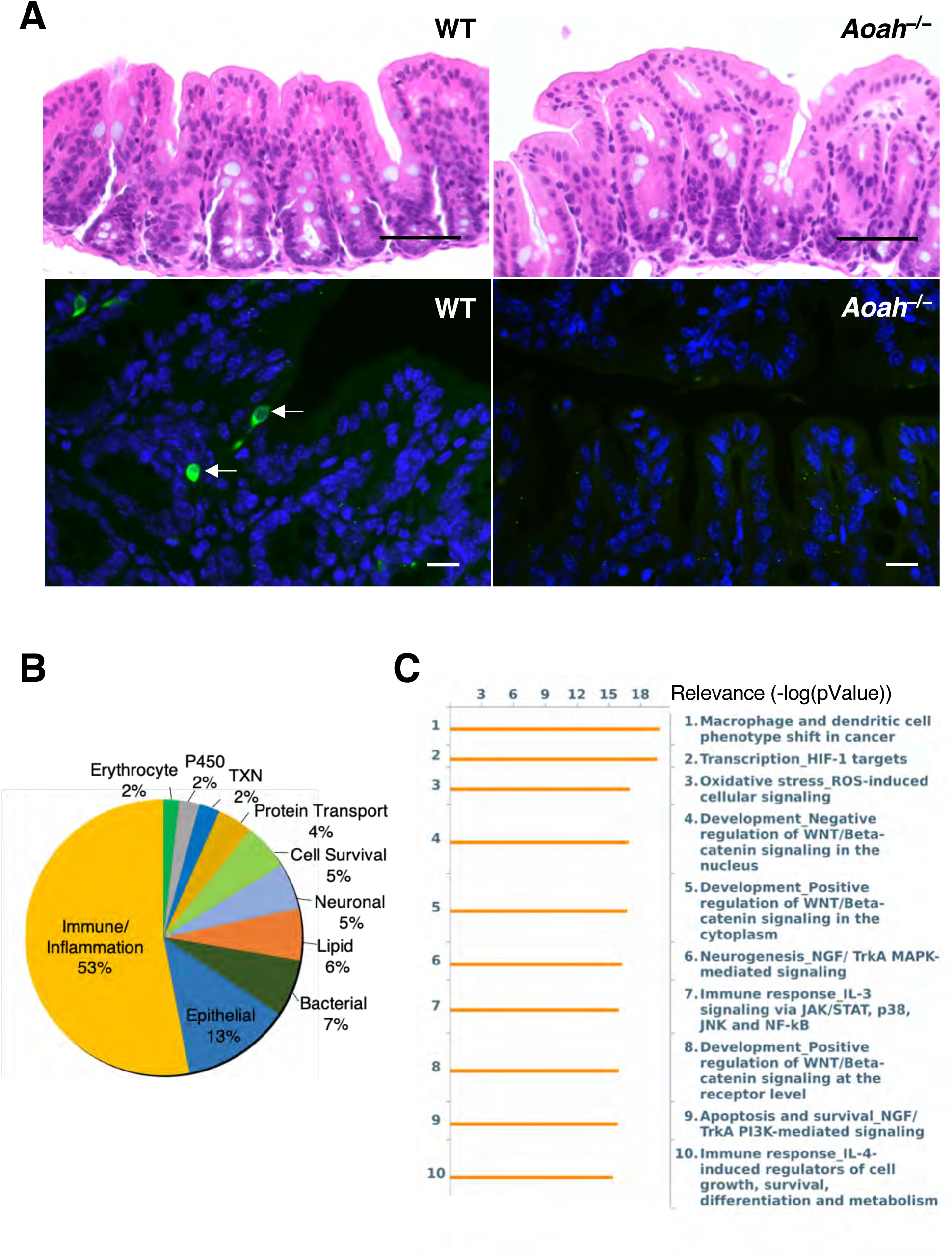
AOAH-deficient mice have normal gut morphology but altered gut signaling pathways. **A**. Comparison of representative cecum H&E staining of female WT (top left) and *Aoah*^—/—^ (top right) mice (scale bar: 50 μm). Immunostaining of AOAH (green) in WT (bottom left) and and *Aoah*^—/—^ (bottom right) cecum. DAPI staining nuclei shown in blue (scale bar: 17 μm). **B**. Pie chart of transcriptome profiling by microarray analysis showing levels of mRNA with 2-fold difference in AOAH-deficient cecum compared to wild-type. **C**. Pathway analysis indicating top 10 signaling pathways altered in AOAH-deficient cecum compared to wild-type.

### AOAH-deficient mice have altered gut microbiota

Previous studies have linked increased gut permeability to gut dysbiosis, and cecal enlargement is a long-known phenotype of germ-free rodents, both suggest potential gut dysbiosis in AOAH-deficient mice [29–31]. We utilized 16S rRNA amplicon sequencing data to assess large compositional differences in fecal (Fig. 3) and cecal (Fig. S3) microbiota between wild type and *Aoah*^—/—^ mice. As shown by principal component analysis (PCA) and the corresponding heat maps, AOAH-deficient mice demonstrate an altered gut microbiome compared to wild type control in both fecal and cecal stool (Fig. 3A and C and Fig. S3A and C respectively). AOAH-deficient mice show group variance along the PC1 axis in bacterial composition of gut flora compared to wild type mice in both fecal and cecal stool (Fig. 3A and S3A, respectively). In addition, α-diversity calculations revealed a greater number of operational taxonomic units (OTUs) in AOAH-deficient fecal stool (444.4±30.67 OTUs) compared to WT fecal stool (366.6±43.93 OTUs, Fig. 3B). We did not observe a significantly different number of observed OTUs between groups in cecal stool (Fig. S3B). However, PC2 axes reveal larger variances amongst individual AOAH-deficient mice compared to wild type, suggesting greater bacterial diversity in mice deficient for AOAH (Fig. 3A and S3A).

**Figure 3.**
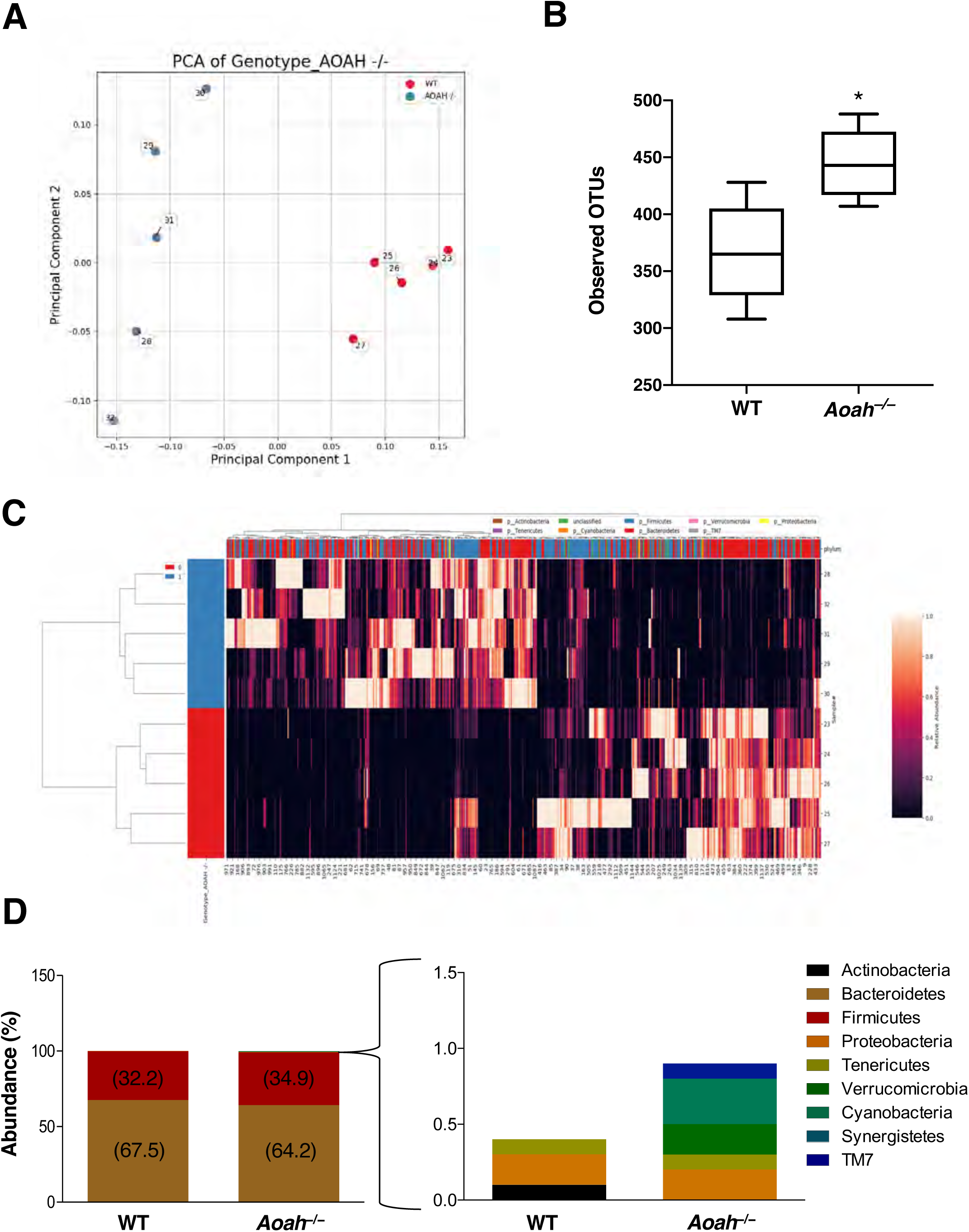
*Aoah*^—/—^ mice exhibit gut dysbiosis. **A**. PCA plot of 16S rRNA analyses of fecal stool. Dots represent individual mice; AOAH-deficient mice shown in blue (n=5) and wild-type mice shown in red (n=5). **B**. α-diversity as measured by observed operational taxonomic units (OTUs), n=5 for both groups, P<0.05. **C**. Heat map of 16S rRNA analyses of AOAH-deficient (blue) and wild-type (red) fecal stool (n=5). **D**. Quantification of bacterial phyla using 16S rRNA sequencing in wild-type and AOAH-deficient fecal stool (n=5).

To determine the specificity of AOAH modulation of gut microbiota, we examined the relative abundance of bacterial phyla (Fig. 3D). Over 99% of bacteria identified in both mouse groups belonged to Bacteroidetes and Firmicutes phyla, with no significant differences in abundance of these groups between wild type and AOAH-deficient stool. However, there were differences in less common phyla such as Verrucomicrobia (0% expression for WT and 0.2% expression for *Aoah*^—/—^), Cyanobacteria (0% expression for wild type and 0.3% expression for *Aoah*^—/—^), and TM7 (0% expression for wild type and 0.1% expression for *Aoah*^—/—^), where AOAH-deficient mice showed more abundant expression of these phyla compared to wild type. A decrease in abundance was observed for Actinobacteria in AOAH-deficient mice compared to wild type (0.10% expression for wild type and 0% expression for *Aoah*^—/—^). These findings suggest a gut microbiome that is richer in diversity in AOAH-deficient mice compared to wild type.

To identify changes in the presence and absence of bacterial species between wild type and AOAH-deficient mice, the top 500 bacterial OTUs were compared in fecal (Table 1) and cecal (Table S1) samples. We observed several bacterial species that were present in AOAH-deficient stool but absent in wild type stool (Table 1). In addition, several bacterial species were absent in AOAH-deficient stool but present in wild type stool. Cecal samples exhibited a modest overlap with our findings in fecal samples but were identified to have several more bacterial species unique to AOAH-deficient mice compared to wild type (Table S1 for full list). Upon further analyses of bacterial functionality, many of the bacterial species with altered abundance were associated with carbohydrate metabolism (Table 1 and Table S1).

**Table 1.** Characterization of Fecal Bacteria in *Aoah*^*—/—*^ Stool Compared to WT

| **Fecal Bacterial Species** | **Absent or Present** | **Functionality** |
| --- | --- | --- |
| *A. onderdonkii* | P | Carbohydrate fermentation; end-product: succinic acid [69] |
| *B. crossotus* | P | Th1/Th2 inflammation balance; fermentation of maltose, starch, glycogen, and dextrin [70, 71] |
| *C. alkalithermophilus* | P | Cellulose and xylan hydrolysis [72] |
| *C. colinum* | P | Fatty acid metabolism; associated with enteric disease [73] |
| *L. longoviformis* | P | Dehydrogenation of enterodiol to enterolactone [74] |
| *L. orale* | P | No active metabolism [75] |
| *O. sitiensis* | P | Diphenol metabolism [76] |
| *P. parvulus* | P | Carbohydrate metabolism [77] |
| *S. aquaeclarae* | P | Production of catalase and gelatinase [78] |
| *A. furcosa* | A | Esculin hydrolysis; end-products: acetic and lactic acids [79] |
| *B. samanii* | A | Glucose, lactose, maltose, sucrose, and trehalose fermentation [80] |
| *C. methylpentosum* | A | Fermentation of methylpentoses and pentoses [81] |
| *L. pectinoschiza* | A | Carbohydrate metabolism [82] |
| *M. plutonius* | A | Glucose, fructose, and mannose metabolism [83] |
| *P. gordonii* | A | L-arabinose production [84] |

### Co-housing alters the functional characterization of bacterial communities

We utilized in silico metagenome prediction of function with PICRUSt to gain functional insights into the effects of AOAH-deficiency and the impacts of microbiome manipulation [26]. PICRUSt revealed that AOAH-deficient mice have several significant differences relative to controls including genes that mediate cell growth and death, signaling molecules and interaction, DNA replication and repair, transcription, human diseases, lipid metabolism, the circulatory system, the digestive system, and the immune system (Fig. 4 and Table S2). Upon further analyses, we observed that co-housing AOAH-deficient mice with wild type mice, to allow for coprophagia-mediated microbiome manipulation, altered the predictive metagenomes in both mouse strains (Fig. 4). Wild type and AOAH-deficient mice showed differences in genes associated with several different taxa, including the RIG-I-like receptor signaling pathway (Fig. 4A) which plays a role in the innate immune system’s ability to recognize pathogens [32]. The differences observed in a subset of functional classes were rescued in co-housed AOAH-deficient mice (Fig. 4A-D and H). Interestingly, for other functional classes, co-housing resulted in no changes in AOAH-deficient mice but altered the predictive values in co-housed wild type to similar values observed in AOAH-deficient mice (Fig. 5E-G). These data suggest that both wild type and AOAH-deficient microbiota can influence changes in the metagenome.

**Figure 4.**
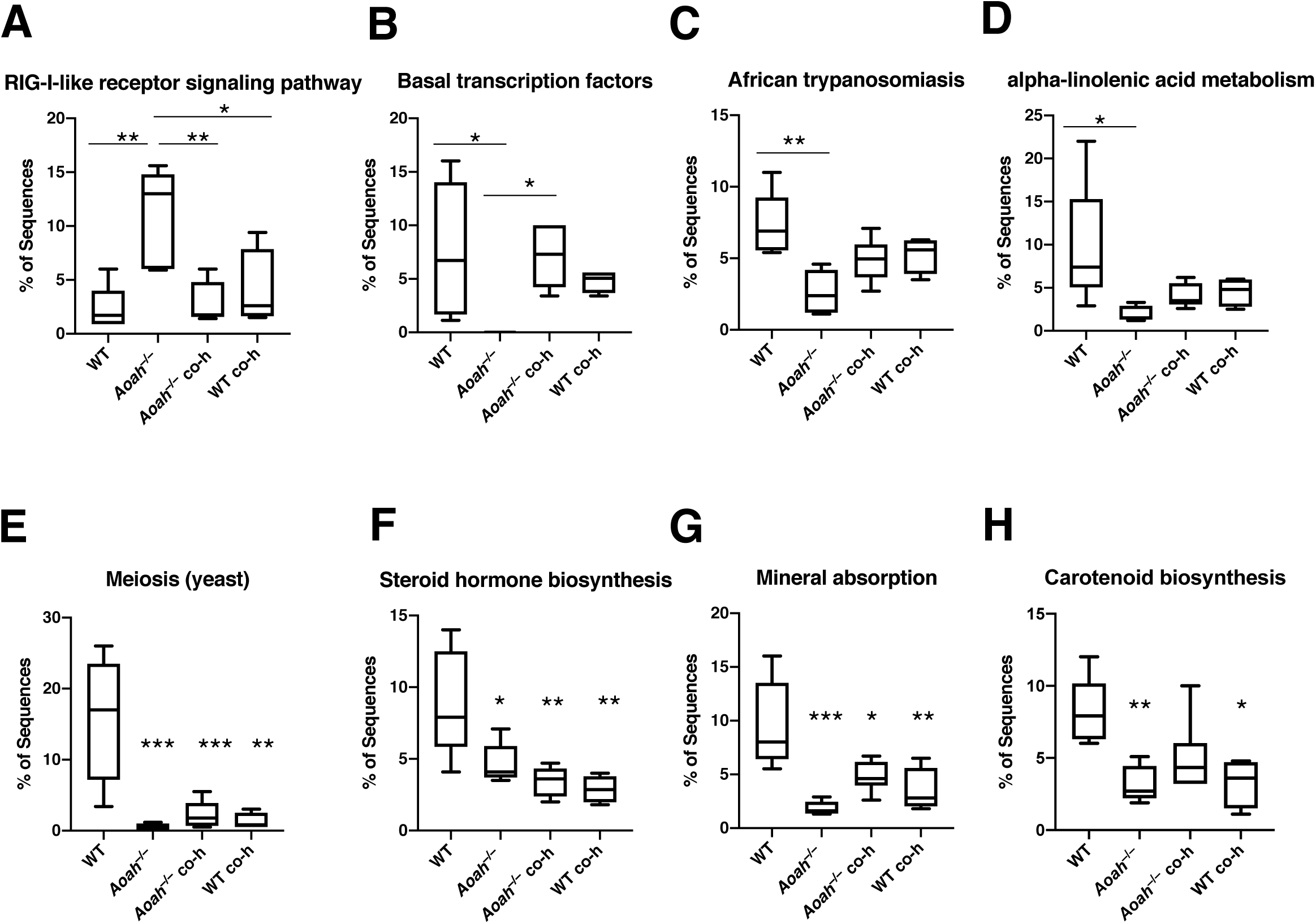
Microbiome-dependent changes in the functional characterization of bacterial communities. Biological functions were predicted by implementing Phylogenetic Investigation of Communities by Reconstruction of Unobserved States (PICRUSt) analyses using 16S rRNA sequence data from wild-type and AOAH-deficient fecal samples. Changes in the gut microbiome by genetic manipulation (*Aoah*^—/—^) or by co-housing (*Aoah*^—/—^ co-h and B6 co-h) resulted in altered predictions including functions related to the RIG-I-like receptor signaling pathway **(A)**, basal transcription factors **(B)**, African trypanosomiasis **(C)**, alpha-linolenic acid metabolism **(D)**, meiosis (yeast, **E**), steroid hormone biosynthesis **(F)**, mineral absorption **(G)**, and carotenoid biosynthesis (H). n=5, *P<0.05, **P<0.01, ***P<0.001, One-Way ANOVA followed by post-hoc Tukey HSD. Data represented as average ± SEM.

**Figure 5.**
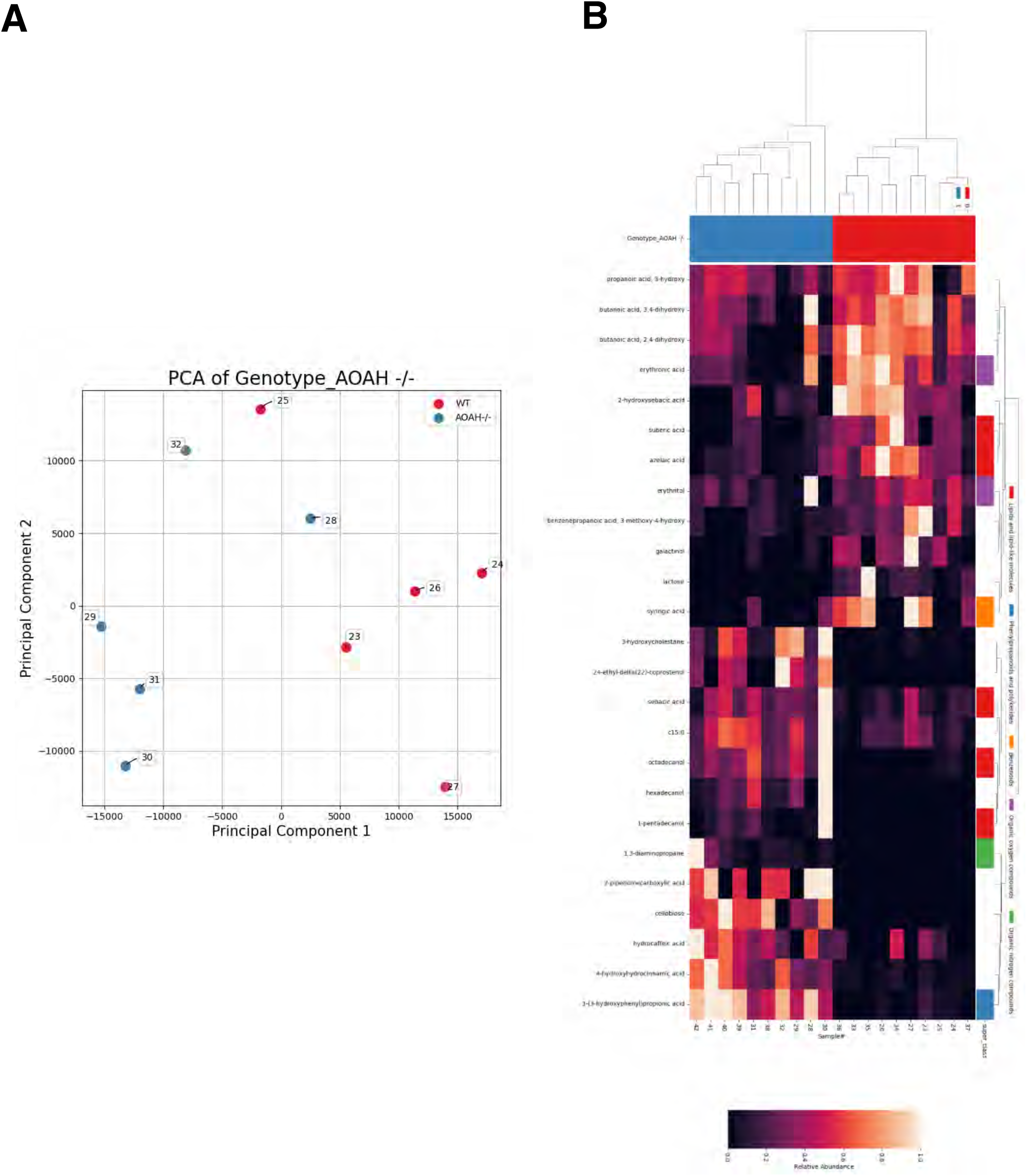
*Aoah*^—/—^ mice exhibit altered fecal metabolomics. **A**. PCA plot of metabolomics of fecal stool. Dots represent individual mice; AOAH-deficient mice shown in blue (n=5) and wild-type mice shown in red (n=5). **B**. Heat map of metabolomics of AOAH-deficient (blue) and wild-type (red) fecal stool (n=5).

### AOAH-deficient mice have altered gut metabolites

To identify the biochemical consequences of dysbiosis associated with AOAH deficiency, we performed untargeted metabolomics. In both fecal and cecal stool, AOAH-deficient mice show altered gut metabolomes compared to controls (Figs. 5 and S3B, respectively). Among the top 500 feature metabolites, we identified numerous metabolites either uniquely present or absent from fecal or cecal AOAH-deficient mice (Tables 2, S3 and S4). We observed that 24-Ethyl-delta(22)-coprostenol, cellobiose, and hexadecanol were present in AOAH-deficient fecal samples but absent in wild type whereas, methionine was absent in AOAH-deficient fecal samples (Table 2). Similarly, cecal samples also expressed the same unique metabolites as fecal samples in AOAH-deficient mice (Tables S3 and S4). In addition to overlap with fecal metabolites, we identified 23 newly present metabolites and 8 absent metabolites in AOAH-deficient in cecal stool compared to controls (Tables S3 and S4). Similar to our observations of bacterial species, functionality of the metabolites with aberrant presence/absence were primarily associated with carbohydrate metabolism (compare Tables 1 and S1 with Tables 2, S3, and S4). In addition, we observed the presence/absence of gut metabolites associated with lipid and amino acid metabolism (Tables 2, S3, and S4). Taken together, our findings show that AOAH-deficient mice have an altered gut microbiome which corresponds with altered accumulation of gut metabolites.

**Table 2.** Characterization of Fecal Metabolites in *Aoah*^*—/—*^ Stool Compared to WT

| Fecal Metabolites | Absent or Present | Functionality |
| --- | --- | --- |
| 24-Ethyl-delta(22)-coprostenol | P | Derived from cholesterol metabolism [85] |
| Cellobiose | P | Reducing sugar; Glucose hydrolysis [86] |
| Hexadecanol | P | Palmitic acid reduction [87] |
| Methionine | A | Polyamine and creatine synthesis; conversion to S-adenosylmethionine; goblet cell development in small intestine [88–90] |

### Co-housing converges microbiota and alters gut phenotype in AOAH-deficient mice

To determine if gut dysbiosis could be rescued in AOAH-deficient mice, we co-housed AOAH-deficient and wild type mice for 7 days to allow for coprophagia-mediated microbiome manipulation. Co-housing resulted in a convergence of cecum size between wild type and AOAH-deficient mice, in contrast with the distinct morphologies in like-housed mice (Fig. 6A, compare with Fig. 1A). We next determined whether our phenotypic observations in co-housed mice would match in genotype. Similar to our observations of converged gut morphology (Fig. 6A), 16S rRNA amplicon sequencing revealed that co-housing resulted in a microbiota genotype that was converged in both mouse strains and distinct from both wild type and AOAH-deficient mice that were like-housed (Fig. 6B).

**Figure 6.**
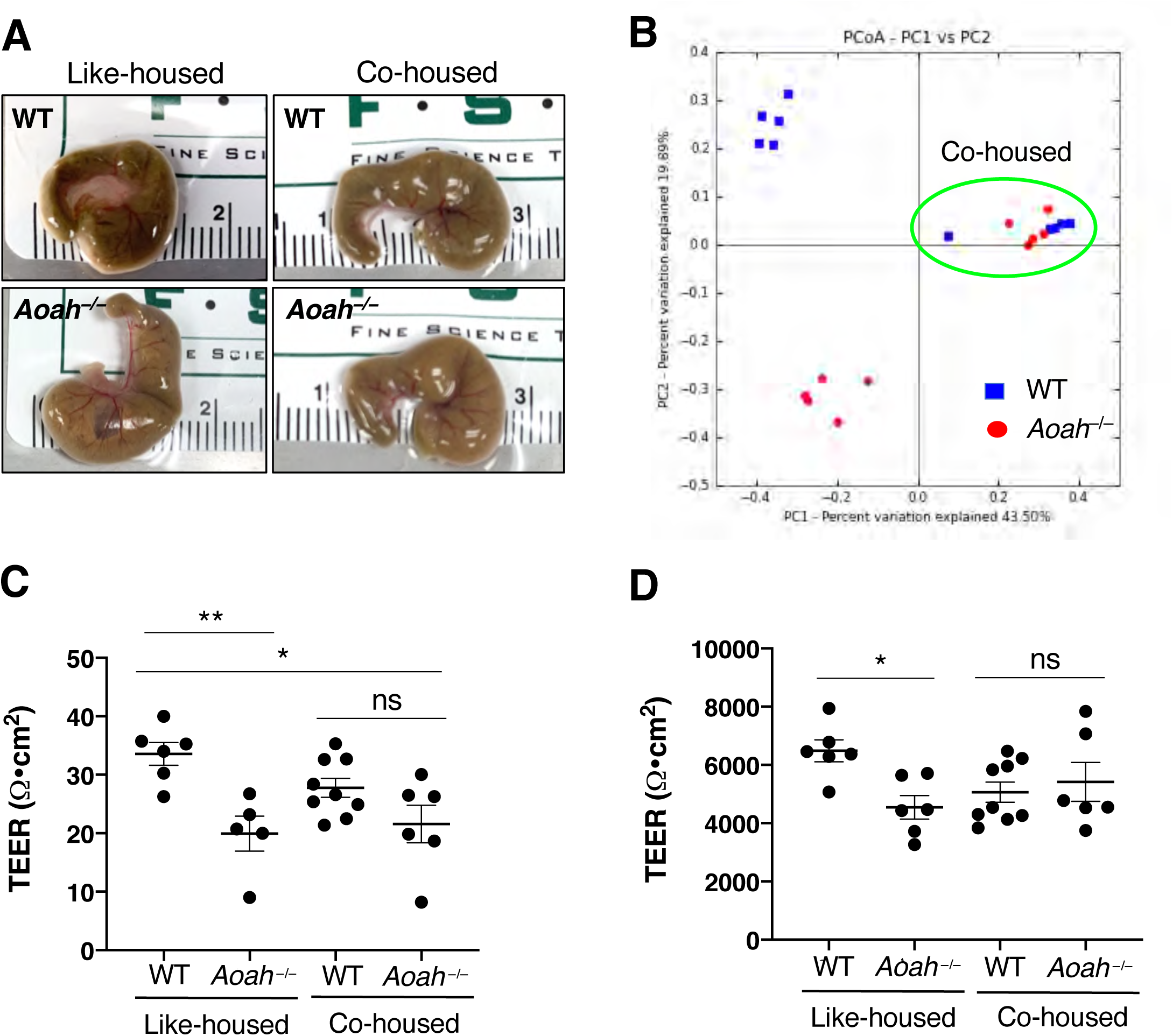
Converging microbiota by co-housing results in altered gut morphology and TEER. **A**. Comparison of representative cecum sizes of female WT (top left), *Aoah*^—/—^ (bottom left), co-housed WT (top right), and co-housed *Aoah*^—/—^ (bottom right) mice. **B**. PCA plot of 16S rRNA analyses of fecal stool in like-housed and co-housed animals. Dots represent individual mice; AOAH-deficient mice shown in red and wild-type mice shown in blue. n=5 for like-housed conditions and n=10 for co-housed condition (circled in green). **C**. *Aoah*^—/—^ mice showed an improvement in cecum TEER after co-housing (like-housed WT mice: n=6, like-housed *Aoah*^—/—^ mice: n=5, co-housed WT mice: n=8, co-housed *Aoah*^—/—^ mice: n=6; *P<0.05, **P<0.01, One-Way ANOVA followed by post-hoc Tukey HSD). Data represented as individual values and average (represented by horizontal line) ± SEM. **D**. *Aoah*^—/—^ mice showed an improvement in bladder TEER after co-housing (like-housed WT mice: n=6, like-housed *Aoah*^—/—^ mice: n=6, co-housed WT mice: n=9, co-housed *Aoah*^—/—^ mice: n=6; *P<0.05, One-Way ANOVA followed by post-hoc Tukey HSD). Data represented as individual values and average (represented by horizontal line) ± SEM.

We next addressed whether co-housing altered the leaky gut phenotype of AOAH-deficient mice (Fig. 6C). Similar to cecum morphology, quantifying TEER showed a convergence of cecum barrier function (4.85 Ω•cm^2^ for co-housed wild type mice, and 21.58 ± 7.85 Ω•cm^2^ for co-housed *Aoah*^*—/—*^ mice; Fig. 6C). However, co-housed AOAH-deficient mice still exhibited significantly increased cecum permeability compared to like-housed wild type mice, suggesting that co-housing does not fully resolve barrier dysfunction in AOAH-deficient mice (Fig. 6C).

IC/BPS patients exhibit defects in bladder barrier function [33–35], and similarly we reported that AOAH-deficient mice have increased bladder permeability. Because previous studies in pelvic organ crosstalk have shown the gut can modulate bladder inflammation and pelvic pain [36–40], we examined whether microbiome manipulation altered the bladder permeability phenotype of AOAH-deficient mice. Co-housed AOAH-deficient mice showed bladder TEER similar to co-housed wild type mice (5062 ± 1037 Ω•cm^2^ for co-housed WT mice and 5416 ± 1634 Ω•cm^2^ for *Aoah*^*—/—*^ mice, Fig. 6D). In addition, bladder permeability was not significantly different between co-housed mice and like-housed wild type mice (Fig. 6D), suggesting that convergence of microbiota can alleviate the leaky bladder phenotype of AOAH-deficient mice.

### Microbiota manipulation alters pelvic pain

AOAH-deficient mice exhibit pelvic pain behaviors that include pelvic allodynia and increased visceromotor response (VMR) [7]. To determine whether altered gut flora affected the pelvic pain phenotype of AOAH-deficient mice, we co-housed wild type and AOAH-deficient mice for 28 d and quantified allodynia in response to von Frey filaments applied to the pelvic region (Fig. 7A). Following co-housing, AOAH-deficient mice showed a 35% reduction in pelvic allodynia from baseline in response to the highest stimulus compared to AOAH-deficient mice prior to being co-housed (Fig. 7A, 83.57 ± 20.98% response at baseline vs 54.00 ± 40.06% response after co-housing). These data suggest that changes in the gut microbiome, in part, can decrease pelvic allodynia in AOAH-deficient mice but do not completely alleviate the pelvic pain phenotype.

**Figure 7.**
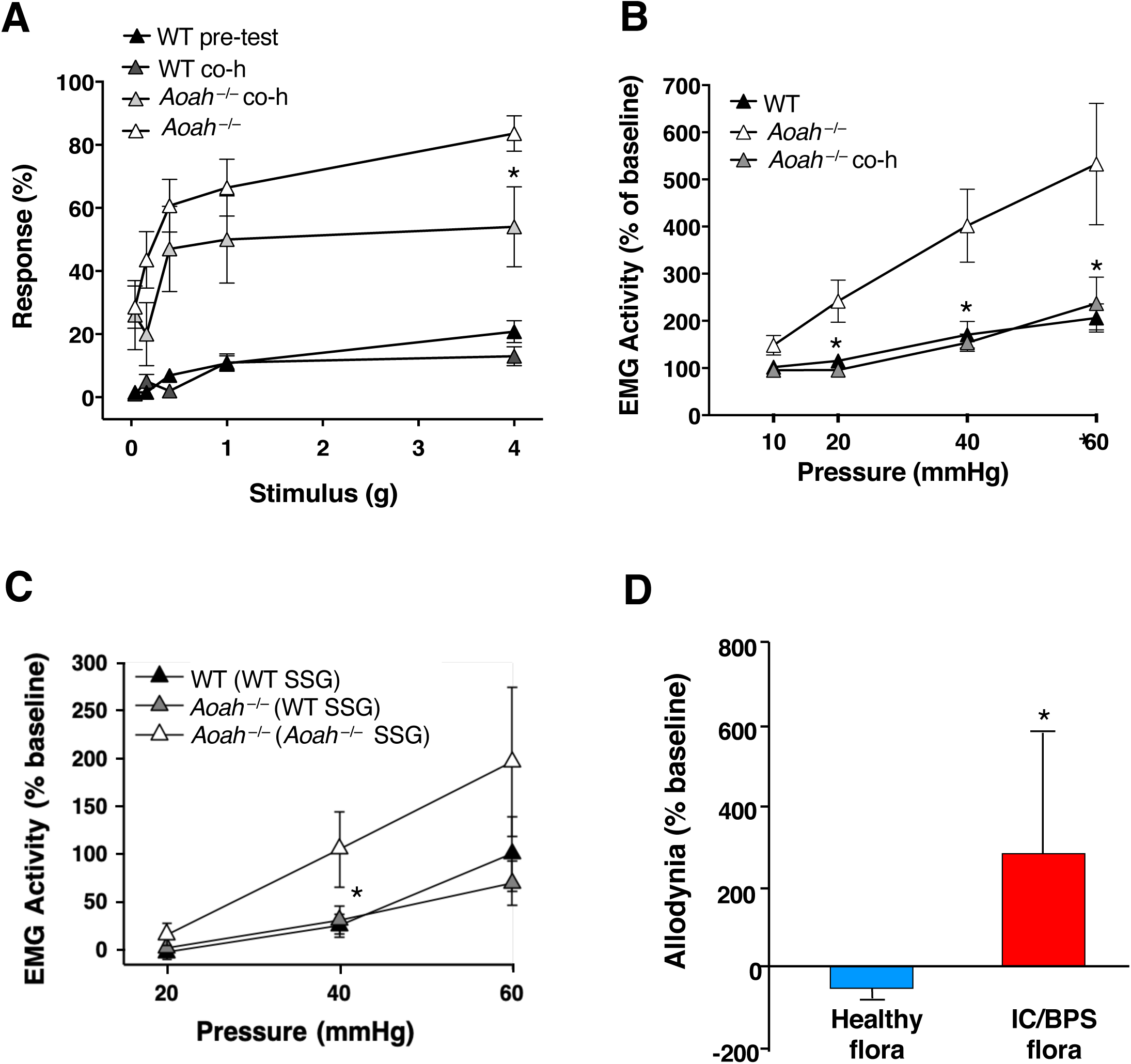
Pelvic pain phenotype is microbiome-dependent. **A**. AOAH-deficient mice exhibited increased pelvic allodynia compared to WT, which was partially alleviated by co-housing for 28 d as shown through response to von Frey filaments stimulating the pelvic region (n=10; *P<0.05, Student’s t test, two tailed). Data represented as average response (%) ± SEM. **B**. Visceromotor response (VMR) was quantified in mice as electromyography (EMG) activity in response to bladder distension under anesthesia. EMG was elevated in AOAH-deficient mice, which was alleviated by co-housing for 7 d (WT mice: n=15, *Aoah*^—/—^ mice: n=9, co-housed (co-h) *Aoah*^—/—^ mice: n=9; *P<0.05, One-Way ANOVA followed by post-hoc Tukey HSD). Data represented as average EMG activity (%) ± SEM. **C**. Measurement of VMR following stool slurry gavage (SSG) from WT and *Aoah*^—/—^ donors. EMG was elevated in AOAH-deficient mice that received SSG from other AOAH-deficient mice compared to WT mice that received SSG from other WT mice. SSG in AOAH-deficient mice from WT donors resulted in reduced EMG activity (WT mice: n=5 and *Aoah*^—/—^ mice conditions: n=4; *P<0.05, One-Way ANOVA followed by post-hoc Tukey HSD). Data represented as average EMG activity (%) ± SEM. **D**. Male AOAH-deficient mice exhibited increased pelvic allodynia following SSG of anaerobic cultures from IC/BPS patients compared to AOAH-deficient mice that received SSG from healthy patients, as shown through response to von Frey filaments stimulating the pelvic region (n=10 for healthy flora, n=12 for IC/BPS flora; *P<0.05, Student’s t test, two tailed). Data represented as average response (%) ± SEM.

To confirm the findings of altered allodynia, we quantified the effects of co-housing on VMR measured by superior oblique abdominal muscle EMG activity following urinary bladder distension (Fig. 7B and C). Co-housing significantly reduced bladder distension-induced VMR in AOAH-deficient mice to levels similar to wild type (Fig. 7B), suggesting decreased pelvic pain following microbiome manipulation. These findings were further corroborated through FMT by stool slurry gavage. AOAH-deficient mice that received gavage of wild type stool slurry showed reduced EMG activity upon bladder distension compared to AOAH-deficient mice that received gavage of stool from other AOAH-deficient donors (Fig. 7C).

Previous reports have shown gut dysbiosis in patients with IC/BPS [19, 20], but whether this dysbiosis modulates pelvic pain remains speculative. To address this question AOAH-deficient mice were gavaged with anaerobic cultures of human stool from healthy controls or IC/BPS patients. After one week of microbiota gavage, mice were assessed for pelvic allodynia. AOAH-deficient mice that were exposed to IC/BPS microbiota showed significantly increased pelvic allodynia from baseline compared to AOAH-deficient mice gavaged with control microbiota (Fig. 7D), suggesting that the gut microbiota of IC/BPS patients can influence pelvic pain. Taken together, our data from manipulating the microbiome of AOAH-deficient mice suggest that gut microbiota modulate pelvic pain.

### Fecal microbiota transplantation reduces anxiety-like behavior in AOAH-deficient mice

We have previously reported that AOAH-deficient mice exhibit comorbid anxious/depressive behaviors, similar to patients with IC/BPS [4, 6, 14]. To assess whether the gut microbiome contributes to the anxious/depressive phenotype of AOAH-deficient mice, we measured defensive burying behavior after placing mice into a novel holding cage with deep, clean bedding. Anxious behaviors were quantified as latency to the first burying event and number of burying behaviors. Corroborating our prior findings, AOAH-deficient mice exhibited higher levels of anxious behaviors as observed by decreased latency to burying and increased number of burying events (Figs. 8A and S4A). Anxiety-like behavior was rescued in AOAH-deficient mice that received gavage of wild type stool, showing longer latency to burying and fewer burying events compared to AOAH-deficient mice that received gavage of AOAH-deficient stool (Fig. 8B and Fig. S4B). Our findings show that the gut microbiome plays a significant role in the anxiety phenotype of AOAH-deficient mice.

**Figure 8.**
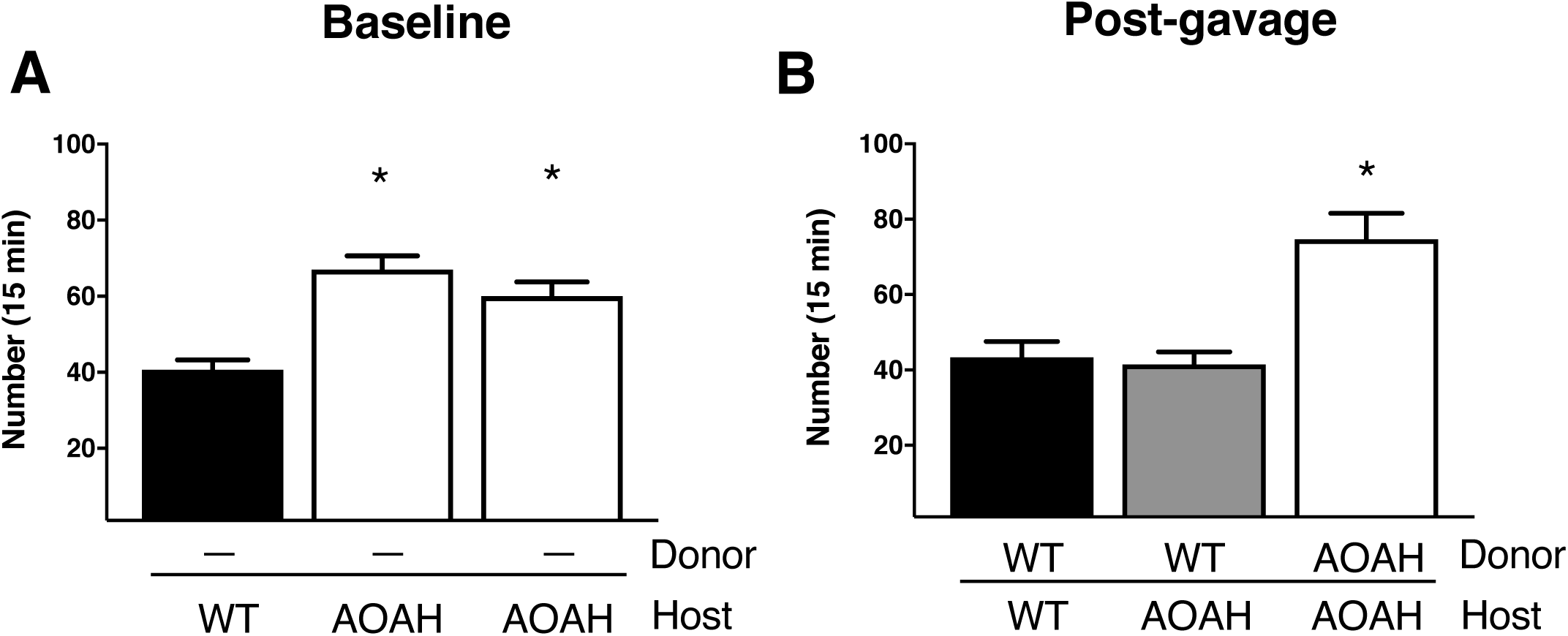
Anxiety-like behavior in AOAH-deficient mice is microbiome-dependent. **A**. Analyses of defensive burying behavior revealed AOAH-deficient mice exhibited increased anxiety-like behavior as measured through number of burying attempts compared to WT mice (WT mice: n=5 and *Aoah*^—/—^ mice conditions: n=4; *P<0.05, One-Way ANOVA followed by post-hoc Tukey HSD). Data represented as average number of burying attempts (for 15 min) ± SEM. **B**. AOAH-deficient mice showed reduced burying attempts 7 d following SSG from WT donors compared to AOAH-deficient mice that received SSG from AOAH-deficient donors (WT mice: n=5 and *Aoah*^—/—^ mice conditions: n=4; *P<0.05, One-Way ANOVA followed by post-hoc Tukey HSD). Data represented as average number of burying attempts (for 15 min) ± SEM.

## DISCUSSION

AOAH deficiency mimics many aspects of IC/BPS, and here we identified *Aoah* as a novel genetic modulator of gut microbiota. Prior studies have implicated host genetic factors in microbiota, including twin studies that showed increased microbiota heritability among monozygotic twins and identified specific host loci in interacting with specific taxa and mediating metabolic processes [41, 42]. Additional human genome-wide association studies (GWAS) and murine quantitative trait loci studies have furthered this understanding of interactions between microbiota and host genetics and provided new mechanistic insights, yet few loci implicated by such studies have been interrogated functionally. For example, 42 human loci were identified that were associated with bacterial diversity, including the vitamin D receptor, and vitamin D receptor-deficient mice had increased Parabacteroides abundance [43]. Although loci on chromosome 7, where *AOAH* resides, were significantly associated with altered microbiota, none of these lay nearer than 2MB from *AOAH*, so GWAS studies have not yet directly implicated human *AOAH* in microbiome modulation. However, *PLA2G3* was significantly associated with microbiota diversity, suggesting a role for arachidonic acid metabolism as a modulator of the human microbiome [43]. That finding dovetails with in silico metagenome analyses that identified arachidonic acid metabolism as significantly associated with microbiota of patients with IC/BPS [19]. We recently reported that AOAH-deficient mice accumulate elevated levels of arachidonic acid and PGE2 in the CNS and found evidence that AOAH is a transacylase mediating arachidonic acid homeostasis via sequestration in phospholipid pools [15]. Thus, together these studies are consistent with an emerging model where loss of AOAH-dependent arachidonic acid homeostasis contributes to gut dysbiosis associated with IC/BPS symptoms.

AOAH-deficient mice exhibited enlarged ceca with decreased barrier function, and transcriptome analyses indicated association with Wnt signaling and inflammatory/immune pathways (Figs. 1, 2 and S1). Altered Wnt signaling in AOAH-deficiency may be due to elevated PGE2 in these mice and the known role of PGE2 as a modulator of Wnt expression [15]. Moreover, Wnt expression has been previously implicated in pelvic pain, where a putative urinary marker of IC/BPS was identified as a glycosylated fragment of Frizzled-8 [44]. Alternatively, in the absence of AOAH that would otherwise detoxify LPS, upregulation of inflammatory and immune responses may be due to excess toll-like receptor (TLR)-mediated signals evoked by LPSs of the gut microbiota. Increased inflammatory responses, in turn, may contribute to loss of barrier function and consequent “leaky gut” [45, 46]. Consistent with this possibility, in silico metagenome analyses by PICRUSt revealed changes in AOAH-deficient mice associated increased RIG-I-like receptor signaling pathway, a regulator of innate immune activation in various cell types [32]. These data were particularly interesting as at least a subset of IC/BPS patients exhibit bladder inflammation associated with activated mast cells [47], and we previously reported that AOAH-deficient mice exhibit significantly more bladder mast cells [7]. In addition, mast cells in the gastrointestinal (GI) tract show increased activation in patients with GI disorders such as irritable bowel syndrome (IBS) [48, 49]. As dysfunctional bowel-bladder crosstalk may also influence GI mast cells in AOAH-deficient mice and gut “leakiness”, activation of these cells should be analyzed.

Immunostaining did not reveal widespread expression of AOAH in gut epithelium or underlying tissue layers (Fig. 2), suggesting that AOAH effects on gut gene expression lie elsewhere. Indeed, we previously reported that AOAH is a genetic regulator of the locus encoding corticotropin-releasing factor (CRF), in brain sites mediating bladder and stress responses where AOAH deficiency leads to increased *Crf* expression and stress responses [13, 14]. CRF alters gut function at multiple levels (reviewed in [50, 51]). At the enteric epithelium, elevated CRF engages CRFR1 receptors on underlying mast cells, resulting in increased epithelial permeability [52]. In addition, centrally administered CRF suppresses gut motility. This raises the possibility that elevated CRF alters cecal motility, contributing to cecal enlargement in AOAH-deficient mice while also altering the lumenal environment and driving dysbiosis. The resulting combination of dysbiosis and leaky gut may then contribute to AOAH-deficient phenotypes of pain and anxiety [17].

Although AOAH expression is not widespread in enteric mucosa, we observed occasional cells within the epithelium that label brightly (Fig. 2). The appearance and prevalence of these AOAH-positive cells in enteric epithelium is reminiscent of neuropods that mediate gut sensory responses [53]. Neuropods are neuroendocrine cells of the enteric epithelium that synapse with vagal neurons, hence providing enteric sensory information to the brainstem through a single synapse and thus poised to convey information about microbiota [54]. Centrally, AOAH deficiency leads to arachidonic acid accumulation, a known mediator of neuronal excitability and nociception [15, 55]. If arachidonic acid metabolism is similarly dysregulated in putative neuropods, this may lead to increased excitability and enhanced sensory responses to microbiota in AOAH-deficient mice. Alternatively, increased intestinal permeability can lead to altered CNS responses when microbial constituents cross the blood brain barrier [46, 56], and future studies will dissect the relative contributions of neuropods and gut leakage to *Aoah* phenotypes.

We observed increased mass of the cecum and cecal stool in male AOAH-deficient mice compared to male WT (Fig. S1), but not in female AOAH-deficient mice compared to female WT (Fig. 1). These findings suggest potential sex differences in cecum phenotype and the differential roles that AOAH may play in cecum development and maintenance. Several studies in mice and humans have shown sex-specific differences in gut microbiota [57]. Although levels of testosterone and estrogen are directly linked to altered gut flora, other factors such as metabolism, body mass index, and colonic transit time differ among males and females and have been linked to changes in the gut microbiome [57], so future studies will examine sex differences in AOAH-mediated microbiota. Nonetheless, we observed significant differences in female microbiota at the levels of phyla and individual taxa, where AOAH-deficient mice showed more bacterial diversity in both fecal and cecal stool and increased verrucomicrobia, cyanobacteria, and TM7 in AOAH-deficient feces (Fig. 3). Cyanobacteria have previously been linked to gastrointestinal symptoms in humans, including abdominal pain, and mucosa of Crohn’s disease and ulcerative colitis patients show significant differences in TM7 bacteria composition [58, 59]. Together these studies suggest a role for cyanobacteria and TM7 phyla in microbiota-associated pelvic pain.

We also observed several OTUs that were present/absent in AOAH-deficient fecal and cecal stool compared to wild type (Table 1 and Table S1 respectively). Based on the known functions of these OTUs, our findings suggest that AOAH-deficient mice may have altered carbohydrate metabolism. This possibility is corroborated by the increased abundance of metabolites that play a role in carbohydrate metabolism (Table 2 and Tables S3 and S4). For example, cecal stool show an absence of *B. acidifaciens* and *P. distasonis* in AOAH-deficient mice (Table S1), bacterial species that regulate glucose metabolism and homeostasis as well as play a role in obesity [60–62]. Previous studies have shown that *B. acidifaciens* and *P. distasonis* prevent obesity or weight gain in mice [61, 62]. Here, we also observed that female AOAH-deficient mice have increased body mass compared to controls, consistent with previous studies that associated reduced *Aoah* expression with increased poultry size [63]. Thus, it is possible that AOAH modulation of microbiota, resulting in altered prevalence of species including *B. acidifaciens* and *P. distasonis* and consequent altered carbohydrate metabolism may explain the increase in body mass observed in diverse species.

Manipulating microbiota demonstrated that gut flora underlie phenotypes of AOAH-deficient mice (Figs. 6-8). Co-housing AOAH-deficient and wild type mice to manipulate microbiota by coprophagia resulted in a merged gut flora composition that was distinct from like-housed mice and corresponded to converged cecum morphology and epithelial barrier function (Fig. 6). In addition, we observed that co-housing or direct FMT by stool slurry gavage resulted in reduced pelvic pain in AOAH-deficient mice (Fig. 7). These findings are similar to the beneficial effects of co-housing on symptomatology and epithelial barrier function of the colon in ulcerative colitis models [64, 65]. However, co-housing colitis mice with controls also increased histologic colitis scores in control mice, suggesting that microbiota transmission may result in a diseased phenotype in previously healthy mice [65]. Although we observed improvement in the pelvic pain phenotype of AOAH-deficient mice after co-housing without adverse pain effects on co-housed control mice, we nonetheless found that co-housing impacted several PICRUSt functional classes in wild type mice, including steroid hormone biosynthesis, mineral absorption, and carotenoid biosynthesis (Figs. 4 and 7). Thus, microbiota alterations associated with AOAH-deficiency are likely sufficient to impact biologic processes and phenotypes.

Our previous studies show that AOAH-deficient mice mimic symptoms and comorbidities of IC/BPS [7, 13, 14]. Here we show that the pelvic pain phenotype and anxiety/depressive-like behavior can be regulated by microbiota (Figs 7 and 8). We identified 2-Quinolinecarboxylic acid, 4,8—dihydroxy (a.k.a., xanthurenic acid) as a gut metabolite of AOAH-deficient cecal stool of mice (Table S3). Xanthurenic acid is a tryptophan metabolite and mGluR_II_ agonist that is elevated in urine of depressed patients, drawing another clinical parallel between patients and AOAH-deficient mice and suggesting this metabolite in AOAH-deficient mice ceca may contribute to the depressive-like phenotype [66–68]. Finally, we found IC/BPS microbiota exacerbate the pelvic pain phenotype of AOAH-deficient mice (Fig. 7). In light of our prior report of altered microbiota among IC/BPS patients [19], this suggests dysbiosis in IC/BPS is functionally significant and sufficient to impact the defining clinical parameter of IC/BPS, pelvic pain. Thus, addressing gut dysbiosis is a therapeutic target for IC/BPS, and novel IC/BPS therapies should be developed that modify the gut microbiome. Such therapies may include prebiotic diets, probiotic supplements to complement deficiencies in specific OTUs or the functions thereof, and even transplant of stool or complex communities derived from stool.

In summary, the data presented here show that AOAH-deficient mice exhibit gut dysbiosis and an altered gut phenotype which mediates pelvic pain and anxiety-like behavior. These findings demonstrate that the gut microbiome is a promising therapeutic target for treating IC/BPS.

## ACKNOWLEDGEMENTS

This work was supported by NIH/NIDDK award R01 DK103769 (B.A.W., A.J.S., and D.J.K) and by NIH/NIDDK T32 DK062716 postdoctoral fellowship to Dr. Rahman-Enyart. Histology services were provided by the Northwestern University Mouse Histology and Phenotyping Laboratory which is supported by NCI P30-CA060553 awarded to the Robert H Lurie Comprehensive Cancer Center. We thank Dr. Robert Munford for generously providing AOAH-deficient mice and for many helpful discussions.

