## Supplemental Data and Tables for "Acyloxyacyl Hydrolase is a Host Determinant of Gut Microbiome-Mediated Pelvic Pain"

**Supplementary Data**

**Figure S1.** Male *Aoah*$^{-/-}$ mice exhibit enlarged ceca, greater mass of cecal contents, and compromised gut permeability. **A.** Comparison of representative cecum sizes of male WT (top) and *Aoah*$^{-/-}$ (bottom) mice. **B.** Male *Aoah*$^{-/-}$ mice showed increased cecal weight versus WT mice (WT mice: n=5 and *Aoah*$^{-/-}$ mice n=4, P<0.001, Student's t test, two tailed) and increased mass of cecal contents (WT mice: n=3 and *Aoah*$^{-/-}$ mice n=4, P<0.0001, Student's t test, two tailed). No differences were observed in body weights (n=4 for both groups, P > 0.05, Student's t test, two tailed). Bars represented as average ± SEM. **C.** Male *Aoah*$^{-/-}$ mice showed a decrease in TEER compared to WT (WT mice: n=9 and *Aoah*$^{-/-}$ mice n=7, P<0.01, Student's t test, two tailed). Data represented as individual values and average (represented by horizontal line) ± SEM.

**Figure S2.** Predictive analysis by transcriptome profiling by microarray analyses indicating top 50 disease risks associated with AOAH deficiency compared to wild-type.

**Figure S3.** *Aoah*$^{-/-}$ mice exhibit cecal gut dysbiosis and altered metabolites. **A.** PCA plot of 16S rRNA analyses of cecal stool. Dots represent individual mice; AOAH-deficient mice shown in orange (n=5) and wild-type mice shown in blue (n=5). **B.** $\alpha$-diversity as measured by observed operational taxonomic units (OTUs), n=5 for both groups, P>0.05. **C.** PCA plot of metabolomics of cecal stool. Dots represent individual mice; AOAH-deficient mice shown in blue (n=5) and wild-type mice shown in red (n=5). **D.** Heat map of 16S rRNA analyses of AOAH-deficient (blue) and wild-type (red) cecal stool (n=5).

**Figure S4.** Microbiome-dependent anxiety-like behavior in AOAH-deficient mice. **A.** Defensive burying behavior revealed AOAH-deficient mice exhibited increased anxiety-like behavior as observed through decreased latency of their first burying attempt compared to WT mice (WT mice: n=5 and *Aoah*$^{-/-}$ mice conditions: n=4; *P<0.05, One-Way ANOVA followed by post-hoc Tukey HSD). Data represented as latency of burying attempt (in sec) ± SEM. **B.** AOAH-deficient mice showed increased latency in their first burying attempt 7 d following SSG from WT donors compared to AOAH-deficient mice that received SSG from AOAH-deficient donors (WT mice: n=5 and *Aoah*$^{-/-}$ mice conditions: n=4; *P<0.05, One-Way ANOVA followed by post-hoc Tukey HSD). Data represented as latency of burying attempt (in sec) ± SEM.

**Table S1.** 16S rRNA sequencing in cecal stool (n=5 for both groups). Represented as bacterial species present (P) or absent (A) in AOAH-deficient cecal stool.

**Table S2.** 16S rRNA sequencing in fecal stool (n=5 for both groups) followed by Phylogenetic Investigation of Communities by Reconstruction of Unobserved States (PICRUSt) to determine predicted functional classes. Data represented as average ± SEM. Student's t test, two tailed.

**Table S3.** Metabolomics of cecal stool (n=5 for both groups). Represented as metabolites present in AOAH-deficient cecal stool.

**Table S4.** Metabolomics of cecal stool (n=5 for both groups). Represented as metabolites absent in AOAH-deficient cecal stool.

**Table S1.** Characterization of Cecal Bacteria in *Aoah*$^{-/-}$ Stool Compared to WT

| Cecal Bacterial Species | Absent or Present | Functionality |
|---|---|---|
| ***A. defectiva*** | P | Carbohydrate fermentation; end-product: lactic acid; risk of causing infective endocarditis [91] |
| ***A. finegoldii*** | P | Carbohydrate fermentation: lactase and succinic acid [92] |
| ***A. massiliensis*** | P | Unknown function [93] |
| ***A. onderdonkii*** | P | Carbohydrate fermentation; end-product: succinic acid [69] |
| ***A. putredinis*** | P | Hydrolyzes tryptophan to indole [93] |
| ***B. cellulosolvens*** | P | Cellulose and cellobiose metabolism; end-product: acetic acid [94] |
| ***B. uniformis*** | P | Carbohydrate metabolism [95] |
| ***C. alkalithermophilus*** | P | Cellulose and xylan hydrolysis [72] |
| ***C. cellulolyticum*** | P | Cellulose catabolism [96] |
| ***C. colinum*** | P | Fatty acid metabolism; associated with enteric disease [73] |
| ***C. thermosuccinogenes*** | P | Fermentation of carbohydrates; end-products: succinate and acetate [97] |
| ***D. russensis*** | P | Sulfur metabolism [98] |
| ***I. melis*** | P | Glucose, glycerol, ribose, and trehalose metabolism [99] |
| ***L. agilis*** | P | Carbohydrate fermentation; End-product: lactic acid [100] |
| ***L. innocua*** | P | β-1,2-Glucan metabolism [101] |
| ***L. orale*** | P | No active metabolism [75] |
| ***M. jinjuensis*** | P | Xylan hydrolysis [102] |
| ***O. sitiensis*** | P | Diphenol metabolism [76] |
| ***P. cellicola*** | P | Glucose fermentation [103] |
| ***B. acidifaciens*** | A | Glucose metabolism (aesculin hydrolysis); prevents obesity and improves insulin sensitivity [60, 62] |
| ***L. pectinoschiza*** | A | Carbohydrate metabolism [82] |

| | | |
|---|---|---|
| ***M. indoligenes*** | A | Glucose, galactose, maltose, and ribose fermentation; End-products: indole, acetate, butyrate, and lactate [104] |
| ***P. distasonis*** | A | Glucose homeostasis; alleviates obesity [61] |

**Table S2.** Predicted Functional Classes Using PICRUSt

| Functional Classes | WT (%) | *Aoah*$^{-/-}$ (%) | p-value |
|---|---|---|---|
| **African trypanosomiasis** | 14.72 ± 4.48 | 5.26 ± 3.10 | 0.0048 |
| **Alpha-linolenic acid metabolism** | 9.80 ± 5.09 | 3.42 ± 1.54 | 0.0455 |
| **Basal transcription factors** | 20.00 ± 16.58 | 0.00 ± 0.00 | 0.027 |
| **Cardiac muscle contraction** | 0.11 ± 0.07 | 19.89 ± 12.21 | 0.0068 |
| **Carotenoid biosynthesis** | 14.36 ± 4.07 | 5.64 ± 2.27 | 0.0030 |
| **Chagas disease (American trypanosomiasis)** | 4.46 ± 4.65 | 15.56 ± 9.12 | 0.0415 |
| **Colorectal cancer** | 0.06 ± 0.09 | 19.94 ± 12.18 | 0.0065 |
| **Ether lipid metabolism** | 20.00 ± 14.34 | 0.27 ± 0.17 | 0.0152 |
| **G-protein coupled receptors** | 19.94 ± 18.56 | 0.03 ± 0.06 | 0.0426 |
| **Influenza A** | 0.06 ± 0.09 | 19.94 ± 12.18 | 0.0065 |
| **Meiosis-yeast** | 19.26 ± 10.93 | 0.66 ± 0.58 | 0.0051 |
| **Mineral absorption** | 16.80 ± 7.48 | 3.20 ± 1.15 | 0.0038 |
| **Non-homologous end-joining** | 15.04 ± 8.11 | 4.96 ± 0.96 | 0.0249 |
| **Parkinson’s disease** | 0.10 ± 0.07 | 19.92 ± 12.21 | 0.0067 |
| **Rig-I-like receptor signaling pathway** | 3.46 ± 3.23 | 16.55 ± 6.95 | 0.0051 |
| **Small cell lung cancer** | 0.06 ± 0.09 | 19.94 ± 12.18 | 0.0065 |
| **Steroid hormone biosynthesis** | 13.19 ± 5.64 | 6.81 ± 2.08 | 0.0447 |
| **Toxoplasmosis** | 0.06 ± 0.09 | 19.94 ± 12.18 | 0.0065 |
| **Viral myocarditis** | 0.06 ± 0.09 | 19.94 ± 12.18 | 0.0065 |

**Table S3.** Cecal Metabolites Present in *Aoah*$^{-/-}$ Stool Compared to WT

| Cecal Metabolites | Functionality |
|---|---|
| **1,3-diaminopropane** | Amino acid metabolism [105] |
| **1-Pentadecanol** | Fatty alcohol; jaspine B synthesis [106] |
| **2-methylsuccinic acid** | Aristolochic acid metabolism [107] |
| **2-oleoylglycerol** | Fatty acid metabolism; oleic acid end product [108] |
| **2-Piperidinecarboxylic acid** | Metabolite of lysine; amino acid metabolism [109] |
| **2-Quinolinecarboxylic acid, 4,8--dihydroxy** | Tryptophan metabolism [110] |
| **24-Ethyl-delta(22)-coprostenol** | Cholesterol metabolism [111] |
| **3-hydroxyphenylacetic acid** | Acetic acid derivative; elevated in urine of autistic children [112, 113] |
| **3-hydroxypyruvic acid** | Serine formation; Derives from glucose metabolism [114] |
| **3-methyl-2-oxobutanoic acid** | Amino acid metabolism; metabolite of valine [115] |
| **Cellobiose** | Reducing sugar; Glucose hydrolysis [86] |
| **Cysteine** | Protein synthesis and generation of sulfur-containing molecules [116] |
| **Desmosterol** | Cholesterol synthesis [117] |
| **Guanosine** | Formation of uric acid [118] |
| **Hexadecanol** | Palmitic acid reduction [87] |
| **Hexanoic acid, 2-hydroxy** | Fatty acid metabolism [119] |
| **Inosine** | Purine metabolism; downregulation associated with depression [120] |
| **Monomethylphosphate** | Metabolite of insecticides [121] |
| **Octadecanol** | Lipid and fatty acid metabolism [122] |
| **P-cresol** | Aromatic amino acid metabolism in intestinal microflora [123] |
| **Putrescine** | Amino acid metabolism [124] |
| **Salicylic acid** | Salicin metabolism [125] |
| **Urocanic acid** | Amino acid metabolism; L-histidine metabolism [126] |

**Table S4.** Cecal Metabolites Absent in *Aoah*$^{-/-}$ Stool Compared to WT

| Cecal Metabolites | Functionality |
|---|---|
| **Butane, 2,3-dihydroxy** | Butene/butanone metabolism [127, 128] |
| **Galactinol** | Galactose metabolism [129] |
| **Galactonic acid** | Galactose metabolism [130] |
| **Hydroquinone** | Benzene metabolism [131] |
| **Pyrophosphate (4:1)** | ATP hydrolysis [132] |
| **Sinapic acid** | Glucose metabolism [133] |
| **Syringic acid** | Metabolism of anthocyanins and other polyphenols in the gut [134, 135] |
| **Trehalose** | Glucose metabolism; carbon/energy source [136] |

**Supplementary Figure 1 Rahman-Enyart et al**

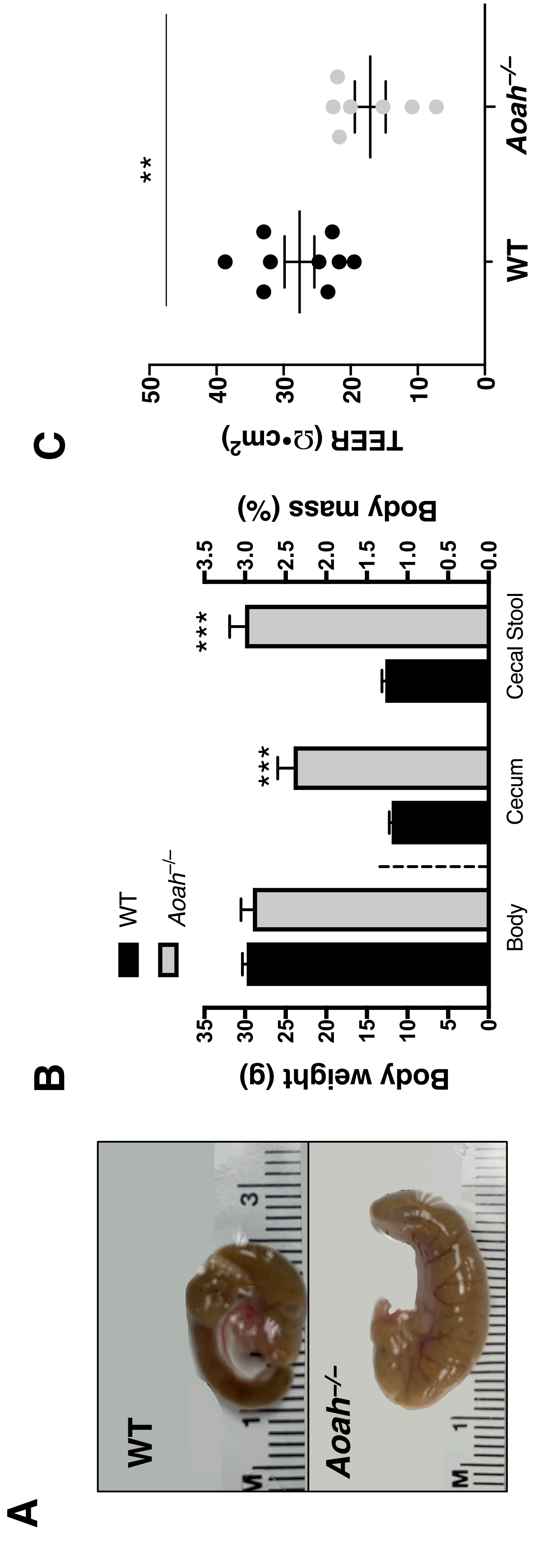

### Supplementary Figure 2 Rahman-Enyart et al.

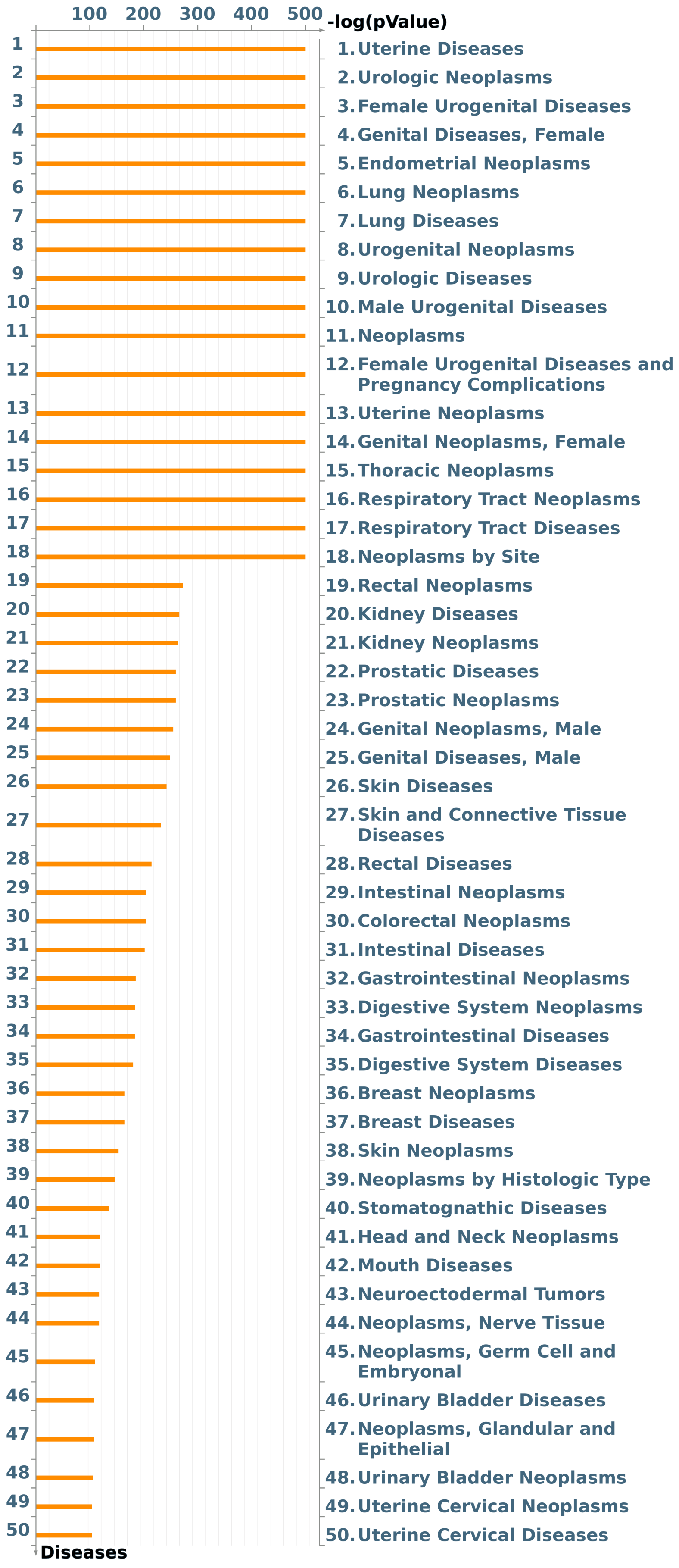

### Supplementary Figure 3 Rahman-Enyart et al.

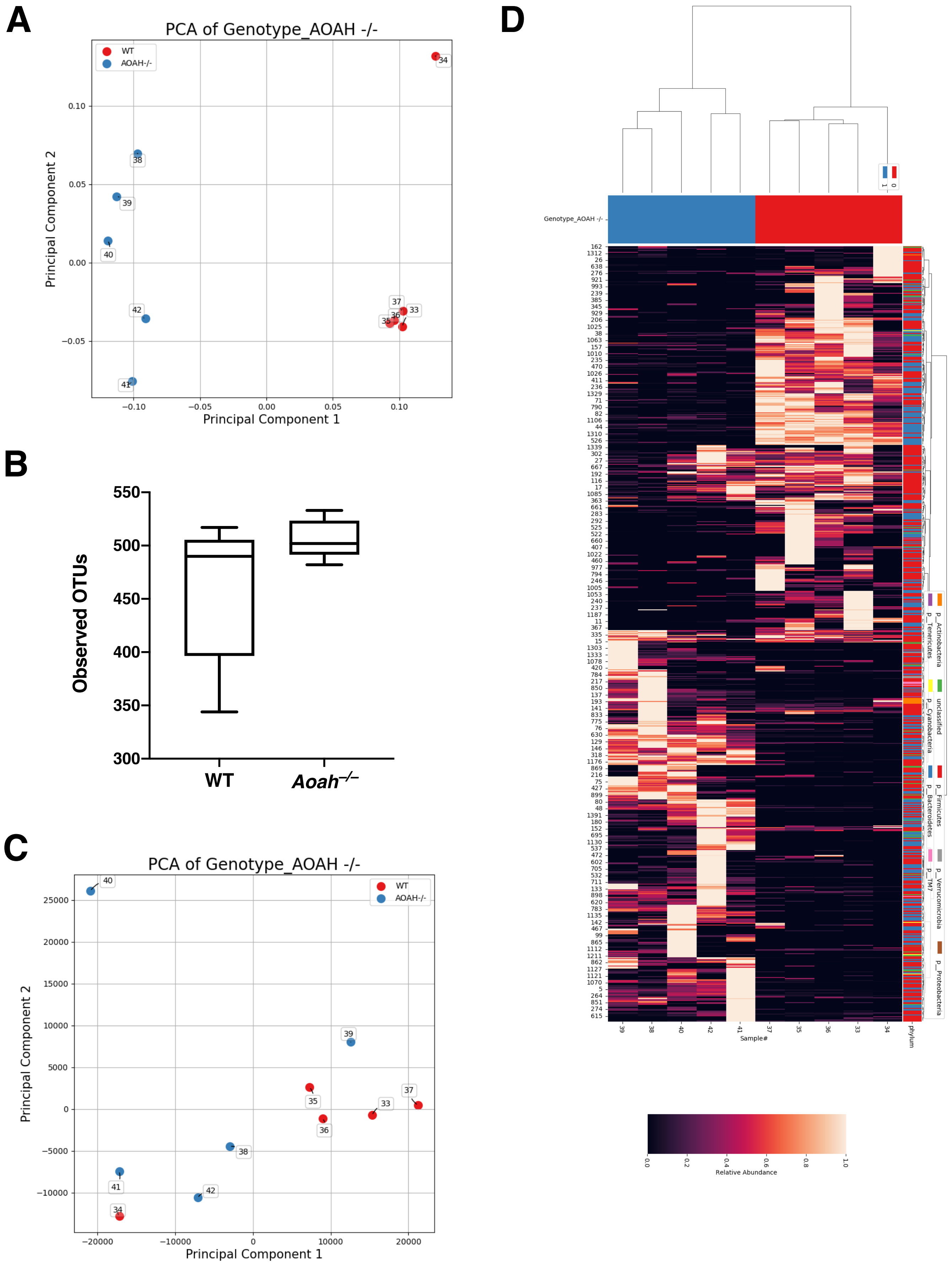

**Supplementary Figure 4 Rahman-Enyart et al.**

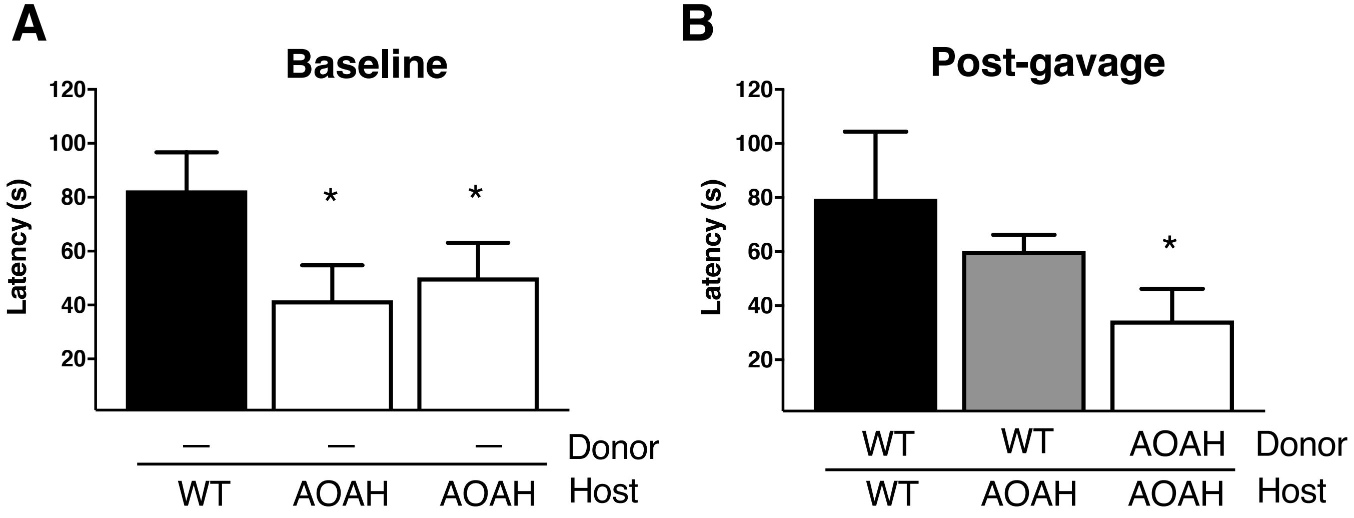
